# Spectral DQE of the Volta Phase Plate

**DOI:** 10.1101/2020.04.20.046110

**Authors:** Bart Buijsse, Piet Trompenaars, Veli Altin, Radostin Danev, Robert M. Glaeser

## Abstract

The Volta Phase Plate (VPP) consists of a heated, thin film that is placed in the same plane as the focused diffraction pattern of an electron microscope. A change in surface potential develops at the point irradiated by the intense, unscattered electron beam, and this altered surface potential produces, in turn, a phase shift between the unscattered and scattered parts of the electron wave. While the VPP thus increases the image contrast for weak-phase objects at low spatial frequencies, we report here that it also leads to the loss of an increasing fraction of the signal at higher resolution. The approximately linear dependence (with increasing resolution) of this loss has been quantified at 200 kV and 300 kV, using evaporated-carbon films of different thicknesses as Volta phase plates. In all cases, the loss of signal remains almost independent of variation of the conditions and parameters that were tested. In spite of having done a number or additional, discovery-based experiments, the cause of this loss of signal remains unexplained at this point.

## INTRODUCTION

The Volta Phase Plate (VPP), introduced by (Danev et al., 2014), provides a significant improvement in contrast for weak phase objects, such as those used in cryo-EM, especially when images are recorded close to focus. This device has been used to produce several groundbreaking results in single-particle cryo-EM (Khoshouei et al., 2017, Liang et al., 2017, Fan et al.,2019). The Volta phase plate has also proven to be of considerable value in electron cryo-tomography (Mahamid et al., 2016, Asano et al., 2015, El Omari et al., 2019, Imhof et al. 2019). In addition, a hole-free phase plate that is similar to the Volta Phase Plate phase plate has also been described (Malac et al., 2012); it has been mainly used in materials science applications (Hayashida et al., 2019).

At the time that the VPP was first introduced, it was stated that the information content in images is reduced slightly due to scattering of electrons that occurs when they pass through the phase plate (Danev et al., 2014). This reduction referred to the relative modulation depth of the contrast-transfer function (CTF), with and without a VPP, and the implicit suggestion was that the loss was constant for all spatial frequencies.

We now report further measurements of the signal-transfer characteristics of the VPP, both at 200 kV and 300 kV, but after optimizing conditions needed to observe the behavior up to 2 Å resolution. In addition to the relatively small, constant offset (reduction) in signal that was noted previously, we now find that the modulation depth for images recorded with the VPP decreases substantially with increasing resolution when compared to that of images recorded without the VPP. Furthermore, the falloff of modulation depth is almost linear with resolution, and the loss occurs even when the defocus value is relatively small. Both of these effects are unlike the nonlinear envelope function due to finite spatial coherence. Although the amount of offset (loss at low resolution) depends, as expected, upon the thickness of the carbon film, the amount by which the signal decreases with increasing resolution is insensitive to the thickness.

A number of additional experiments were thus performed in order to distinguish whether the loss of signal could be attributed to electron interactions that occur either at the surface or within the body of the VPP. The results obtained make it unlikely that interactions within the body of the VPP cause the observed signal loss. Still remaining, therefore, is the possibility that some type of surface interaction, perhaps due to rapid fluctuation of the Volta potential itself as images are being recorded, may be responsible. However, this latter possibility has also been made difficult to envision by results obtained when comparing the performance of the VPP with that of a hole-based phase plate, where – in an ideal case – the Volta potential does not play any role. Perhaps surprisingly, the same resolution-dependent loss of signal occurred when the unscattered beam passed through a relatively large, 15 µm diameter hole in the carbon film. Additional, discovery-based experiments are described in the Results, but the physical cause of the resolution-dependent loss of signal remains unexplained at this stage.

## MATERIALS AND METHODS

Experiments were conducted on various Titan Krios systems, equipped with Falcon cameras and Volta Phase Plates. The Krios AutoCTF software was used for setting the desired defocus and for correction of 2-fold astigmatism. Positions on these phase plates were used only if the Thon rings became round after alignment and correction of 2-fold astigmatism. Obtaining round Thon rings is not always achievable for all areas of a VPP, presumably due to variations in surface topography (wrinkling) or the presence of contaminated areas producing local charging of a VPP.

The Falcon cameras were used in integrating mode, with a typical exposure at the specimen of >100 e-/pix/s and integration times of a few seconds. The pixel sizes, referred to the specimen, varied between 0.5 and 1Å. The 4k x 4k images were analyzed in 2k x 2k sub-areas to minimize effects on the modulation depth of the CTF due to variations in specimen-height over a large field of view.

The specimen consisted of a carbon film, less than 3 nm thick, supported on lacey carbon support film grids (Product No. 01824,Ted Pella). Thin carbon films are required for Thon rings to extend to resolutions as high as 2 Å. This is because the modulation depth decreases at high resolution, while the midpoint of their oscillation does not change, when thick carbon films are used. These specimen-thickness effects are due to curvature of the Ewald sphere, as is pointed out in (DeRosier et al., 2000).

The specimen was cooled to liquid-nitrogen temperature, which is the normal situation in Krios microscopes. The purpose in this case, however, was to ensure that hydrocarbon contamination did not build up in the illuminated area, something that is difficult to avoid for samples imaged at room temperature.

The 2-D power spectra of images were computed using a home-built MatLab script and fitting routines offered by MatLab. The corresponding noise-only power-spectra were obtained by recording images without a sample, both with and without a phase plate. From the 2-D power spectra, radial averages were normally produced by integrating over the full, 0 to 180 degrees azimuthal range. If some remaining 2-fold astigmatism was detected, however, the integration was limited to a smaller azimuthal range, and the results from different azimuthal wedges were separately processed and averaged for the end result. Anisotropic magnification effects were of the order of 1%, which is small enough to ignore in the analysis.

## RESULTS

Radial averages of 2-D power spectra were used to compare the intensities of Thon rings, i.e. the CTF-modulated signal strength, for images recorded with and without a Volta phase plate. All other instrumental parameters were held constant when making this comparison. As is shown in by the example Figure 1, the reduction in the high-resolution, CTF-modulated component of the power spectrum of images that were recorded with a Volta phase plate is already apparent by eye, even before further processing and correction.

**Figure 1.**
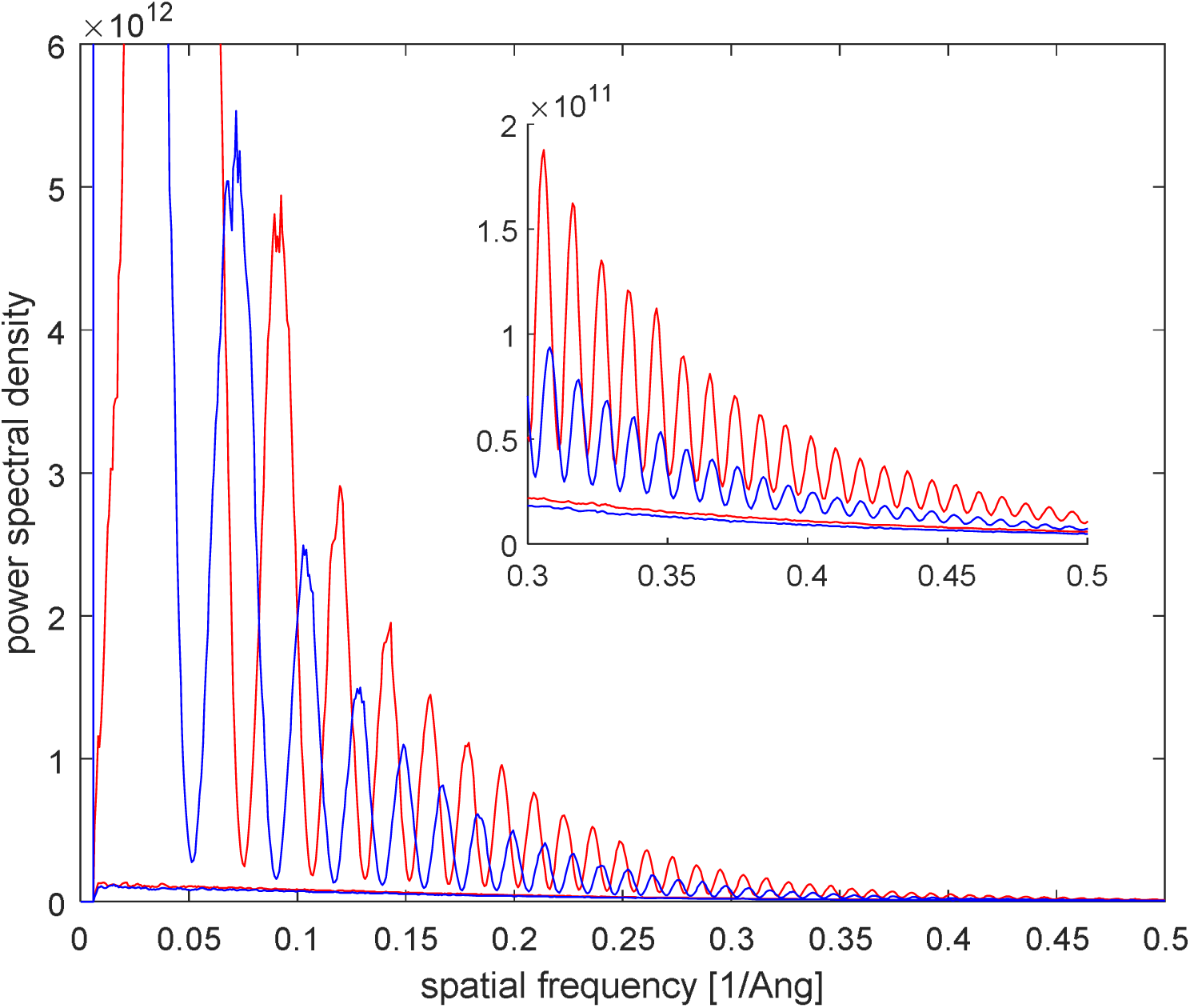
Power spectral density (PSD) of 300 kV images of thin amorphous carbon obtained with a VPP (blue) and without a VPP (red), using an image pixel size of 0.67Å (referred to the specimen) and 2k x 2k pixels. The flat curves are taken without a specimen (“background” curves). The inset zooms in on the high-resolution region. The VPP clearly introduces a reduction of modulation depth of the PSD. This VPP had an estimated thickness of 15-20 nm.

Quantitative comparison, however, requires that one should first correct for the small offset (reduction) in the modulation depth of the Thon rings that is due to the loss of some electrons as they pass through the carbon-film phase plate. The resulting loss of electrons was measured by recording images without a specimen, and comparing the average intensity per pixel, as well as the noise spectral density, for images obtained with and without VPP. The resulting loss of electrons can be seen in the zoomed-in insert (high resolution portion) of Figure 1. This reduction is independent of resolution, as is shown in Figure 2. To correct for this systematic difference, raw power spectra of images were scaled by an appropriate, constant factor, an example of which is shown in Figure 2.

**Figure 2.**
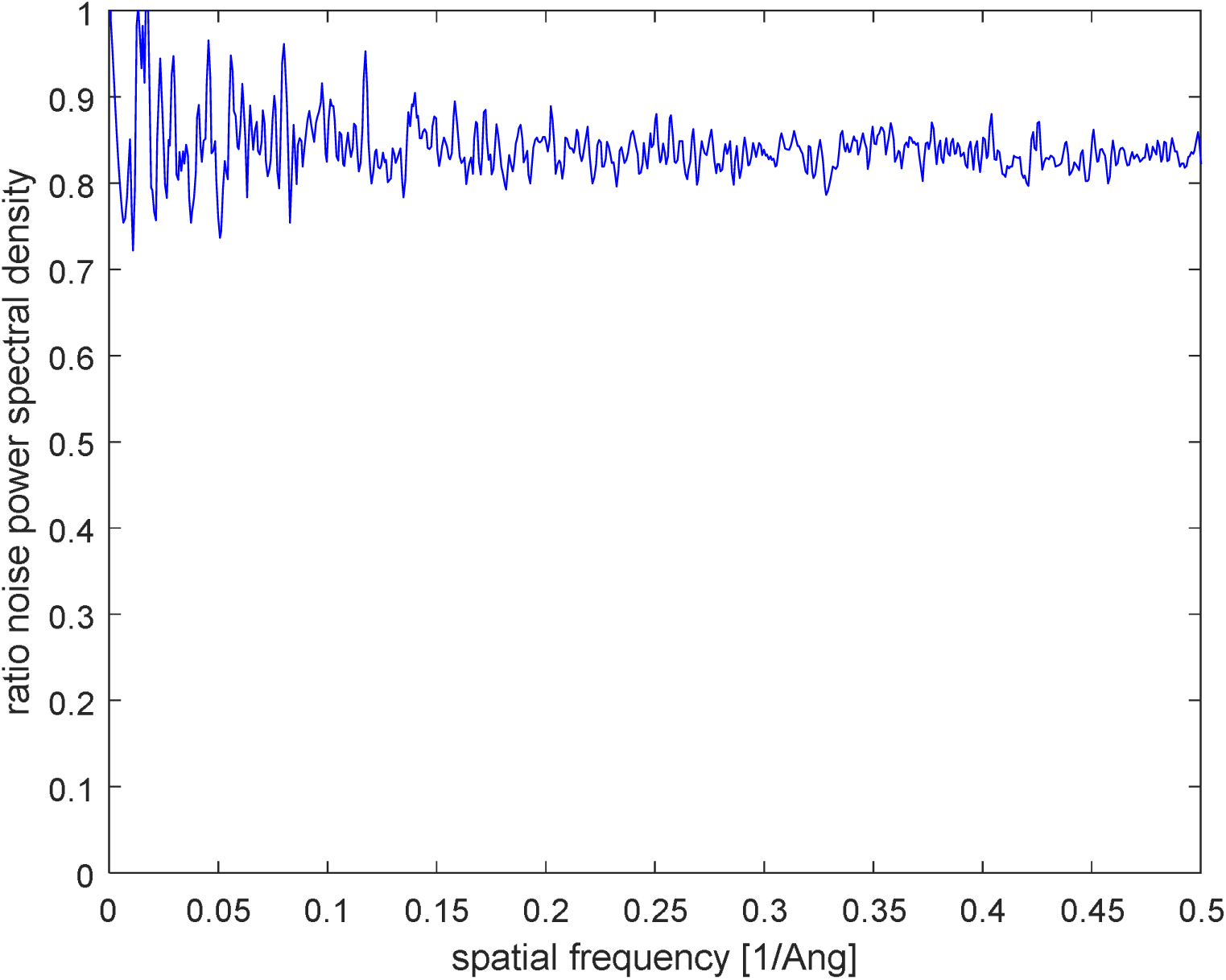
The ratio of the power spectra of background curves as shown in Figure 1 (VPP divided by no VPP). This ratio is constant and has an average value of 0.84.

In addition, one must account for the fact that the local minima of the power spectra, whose positions correspond to zeros in the CTF, do not go to the “zero baseline” associated with electron shotnoise in the power spectra. The gap between the two may be due, at least in part, to non-linear terms in the image intensity. These terms include dark-field contributions to the images of the carbon film that are produced by inelastically scattered electrons as well as by elastically scattered electrons. These contributions are usually ignored when discussing the theory of image formation for a weak phase object. Further investigation of factors responsible for the gap were thus judged to be outside the scope of the main issue in the current study.

In order to eliminate the effect of any differences in the baseline, the upper and lower bounds of the CTF modulation itself were estimated by fitting spline curves to the values at the maxima and minima, respectively, of the oscillations in the power spectra. An example of the resulting spline curves, fitted to these guide points, is shown in Figure 3. The distance between the upper and lower curves, at a given spatial frequency, was then used to estimate the resolution-dependent fall-off of the modulation depth.

**Figure 3.**
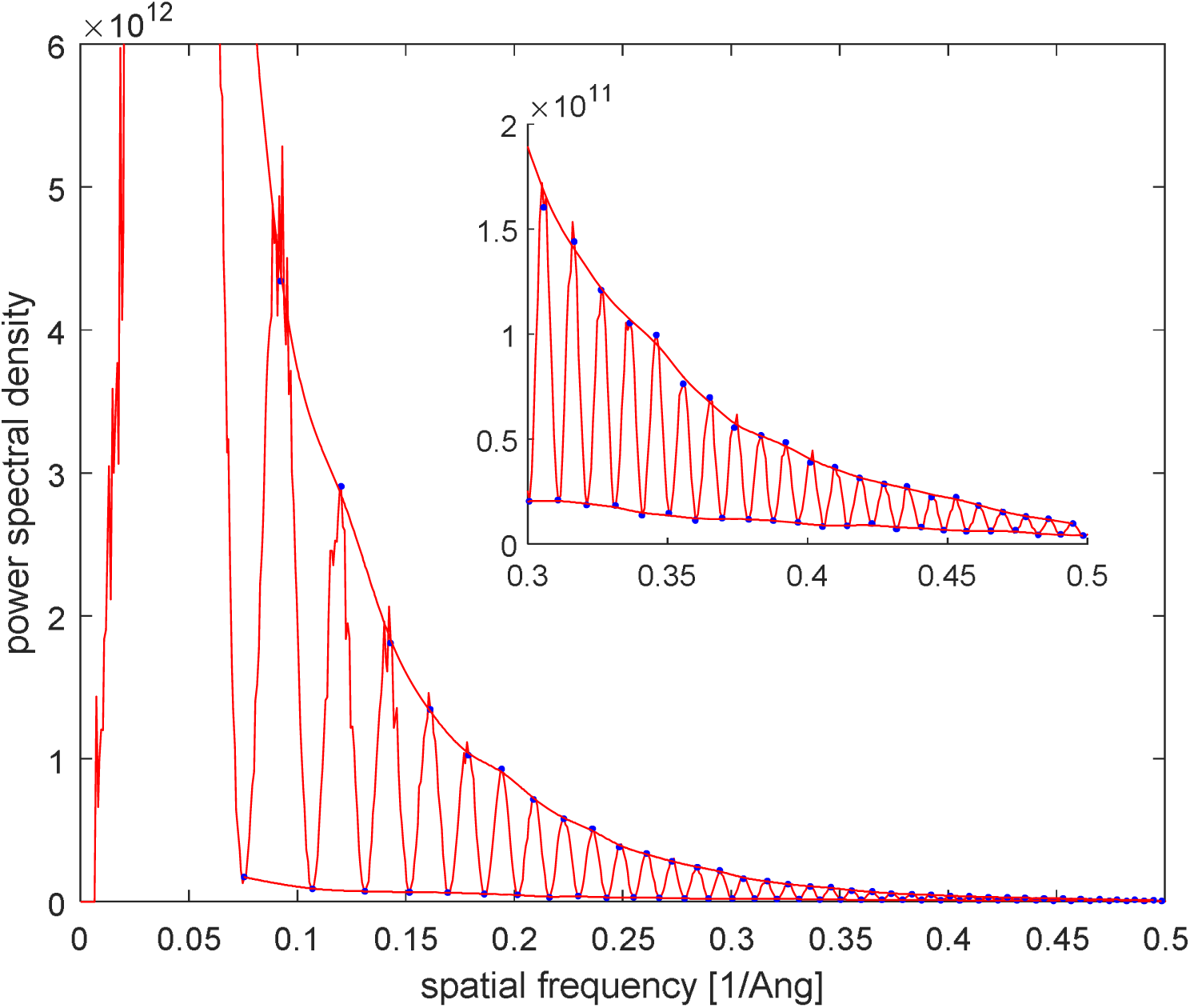
Example of a spline-fit through minima and maxima of a PSD graph.

Finally, the ratio of the two modulation depths, i.e. with and without using a VPP, provides the desired, quantitative comparison of the signal strength for the two cases. Factors responsible for a fall-off of the power spectra that are common to both cases, such as the MTF of the camera, the structure factor of the specimen, and the spatial and temporal coherence envelopes of the microscope, are thereby eliminated. Any remaining fall-off that is seen in the ratio must be due solely to the VPP. We thus define the ratio to be the Detective Quantum Efficiency, DQE_VPP_(s) of the Volta phase plate. Note that DQE is usually defined as the ratio of squared signal-to-noise-ratio (SNR). Because we normalized all our results to the same electron exposure, however, it suffices in our case to take the ratio of the modulation depths of the power spectra.

Figure 4 shows an example, which is based on the raw data shown in Figure 1, of DQE_VPP_(s) for 300 kV electrons. The phase plate in this case consisted of an evaporated-carbon film estimated to be in the 15-20 nm thickness range. The DQE value is already reduced by a factor of ∼0.85 at a resolution of ∼2 nm, even though the data were previously corrected for the loss of electrons that occurs as they pass through the phase plate. The DQE then continues to fall with increasing resolution, becoming ∼0.4 or less at a resolution of ∼2 Å. This experiment was repeated several times, and in all cases the extrapolated value of DQE_VPP_ (0.0 Å^−1^) was about 0.9, and the estimated value of DQE_VPP_ (0.5 Å^−1^) was about 0.35, both with an estimated error of about ±0.05.

**Figure 4.**
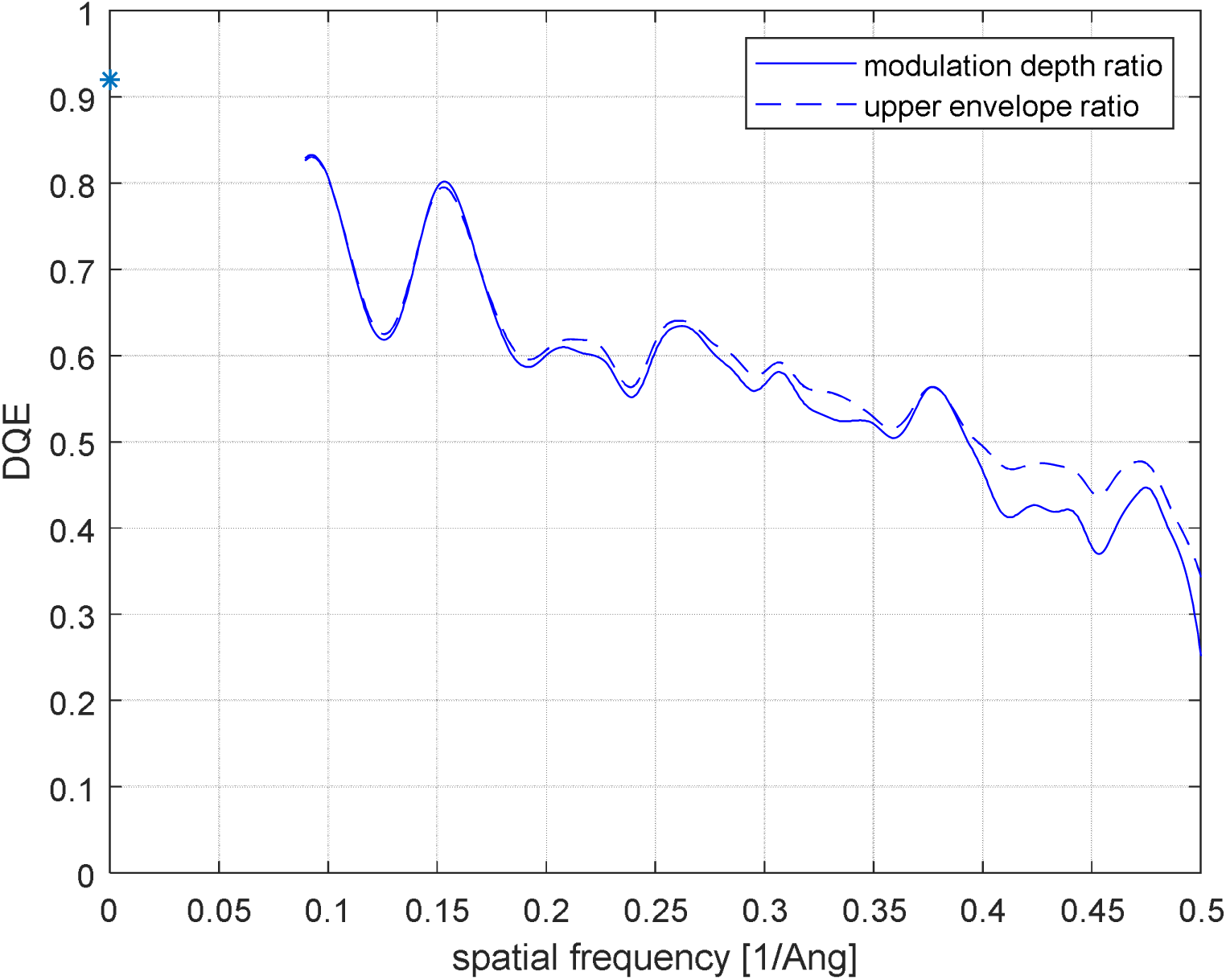
Spectral DQE of the VPP, defined as the ratio of modulation depth of the PSD with and without a VPP. The PSD graphs have been scaled to the same noise spectral density as defined by the background curves. For comparison, the ratio of the upper-envelope spline fits is also shown, and this ratio demonstrates the same behavior as does the ratio of modulation depths. Beam voltage was 300 kV and the VPP thickness was 15-20 nm.

The modulation fall-off is predominantly caused by the reduction in the heights of the CTF maxima, as opposed to a rising value at the minima of the CTF, as is also shown in Figure 3. While the fall-off thus is similar, in this respect, to the fall-off caused by imperfect spatial coherence of the incident electrons, the effect of the latter – if any – would have to be the same with and without a VPP. In addition, we found – as was expected – that the measured DQE did not depend upon the defocus value used to record the images, in the range explored between 50 nm and 500 nm under-focus.

This observed fall-off appears to be independent of carbon-film thickness, within experimental error. For example, Figure 5, shows the result obtained with a carbon-film VPP that was estimated to be only 5 nm thick. In addition, DQE values of a given Volta phase plate were found to be only marginally smaller at 200 kV than at 300 kV, i.e. the difference again appeared to be perhaps within experimental error for what was observed at 300 keV (data not shown).

**Figure 5.**
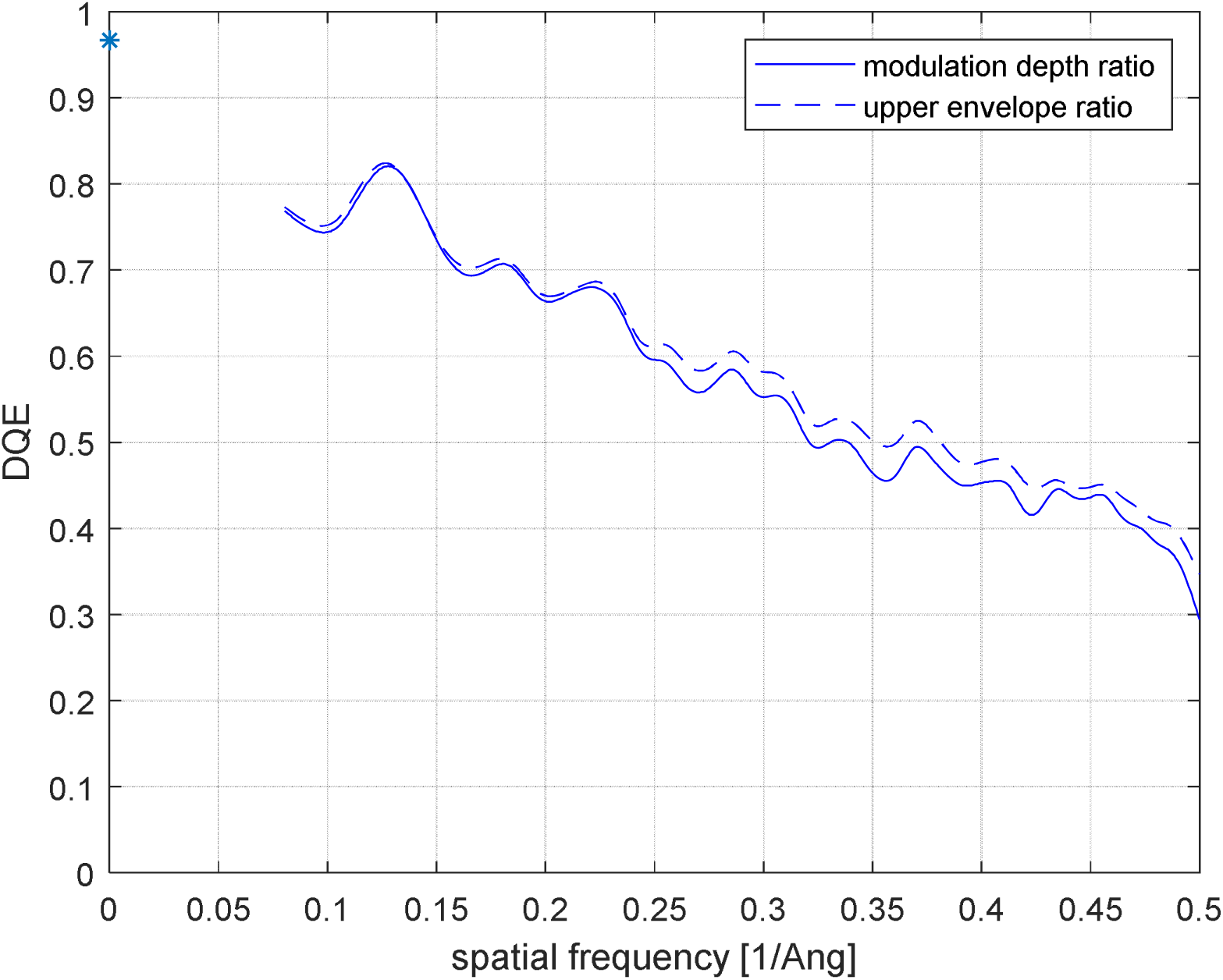
Similar DQE graph as in Figure 4 for a VPP with an estimated thickness of 5 nm, again at 300 kV. The DQE curve fall-off is the same as for a VPP with standard thickness value of 15-20 nm.

The DQE values were unaffected when the electron diffraction pattern was not focused exactly at the plane of the phase plate (off-plane condition), resulting in the unscattered electrons being spread over a larger spot on the carbon film. Even when the spot was intentionally spread to a diameter of 10 µm, the reduction in modulation depth still remained the same, within experimental error.

The reduction in modulation depth at ∼2 Å resolution is similar to what is expected to occur if the energy spread of the electrons were increased by an additional ∼1 eV on passage of the electrons through the phase plate. Even though the shape of the fall-off with increasing resolution is not at all similar to that of a temporal-coherence envelope function, we still considered it important to determine whether an additional, ∼1 eV energy spread really happens.

The energy spread was therefore measured in a Titan microscope operating at 300 kV and equipped with a monochromator and Gatan Imaging Filter. A phase plate was loaded as specimen, using a heating holder to bring the phase plate to a temperature of 250 °C. A Volta spot was created on the phase plate using a beam with current in the range 0.1-0.2 nA. The width of the zero-loss peak was measured without and with sample present. Figure 6 shows that the full width at half maximum of the energy distribution of the unscattered electron beam remained unchanged, as was expected, when a Volta phase plate was inserted. In addition, the energy spread of 0.88 eV is the value usually expected for an FEG-based microscope. Taking the point even further, the energy spectrum of electrons transmitted through a Volta phase plate was measured to be as narrow as 0.11 eV when a gun monochromator was used. This demonstrates that the Volta phase plate does not produce a significant increase in the energy spread of the transmitted electrons.

**Figure 6.**
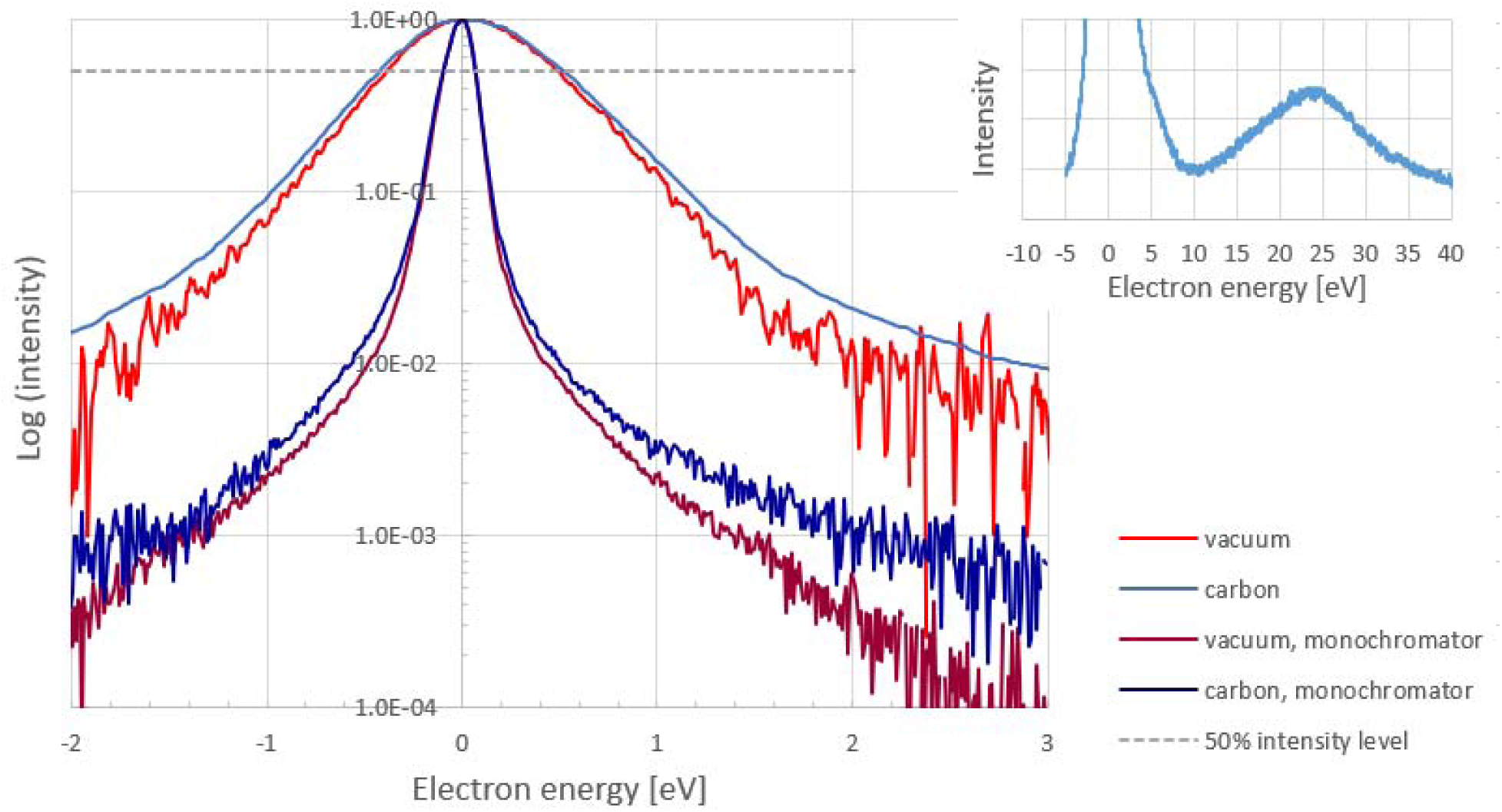
Comparison of EELS spectra obtained either with or without a VPP, inserted as a sample into a Titan column (FEG, 300 kV) fitted with an energy filter (Gatan), and heated to 250 ºC. A Volta potential was created and the zero-loss peak width was recorded, both with (lower, more narrow, dark blue spectrum) and without using a gun monochromator (upper, broader, light blue spectrum). The spectrum extending out to a loss of 40 eV, in the latter case, is inserted in the upper right-hand corner of the figure. As is shown by the red curves, no significant change in the full width at half maximum of the zero-loss peak was observed when moving the sample in and out. All spectra are normalized to a value of 1.0 at zero energy loss. The dashed grey line shows the 50% intensity level from which the full width at half maximum can be judged.

In order to better understand the root cause of the DQE fall-off, we have done two versions of an additional experiment that used a phase plate into which ∼15 µm diameter holes were drilled with a focused ion beam. In the first version, the central, i.e. unscattered, electron beam was focused at the center of one of the holes, and we observed, as is shown in Figure 7, that the DQE remained similar to the situation when a continuous (hole-free) carbon film was used. In the second version, shown in Figure 8, we placed the central beam on the carbon film between two holes. In this way we could determine the DQE fall-off over the spatial frequency band for which the scattered electrons passed through the hole rather than through a carbon film. Together, the results of this experiment demonstrate that the DQE fall-off is identical, within our experimental error, when the unscattered electron beam, the scattered-electron beam, or both pass through a thin carbon film.

**Figure 7.**
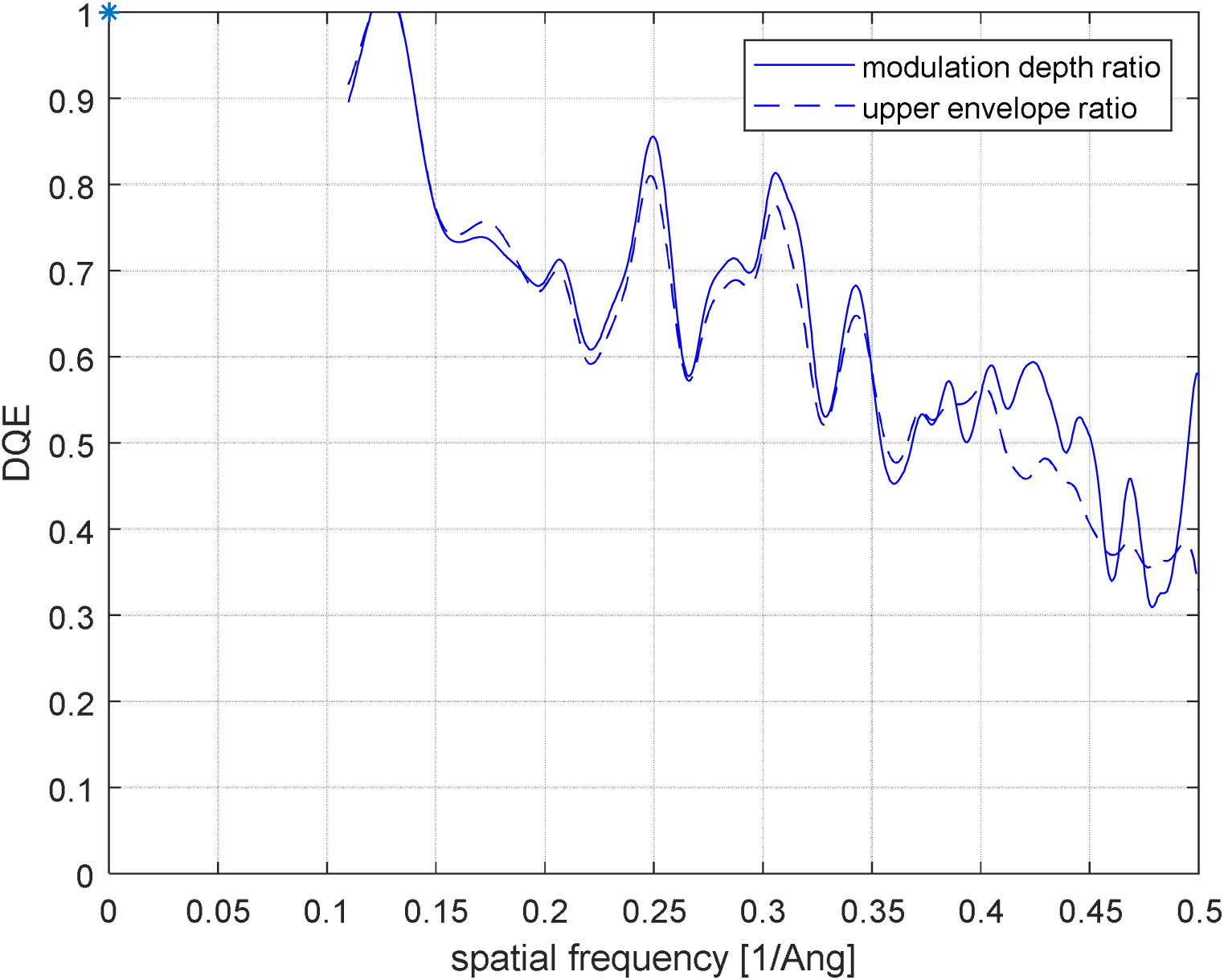
Spectral DQE curve obtained with a ∼3 nm thin amorphous carbon film serving as a hole-based phase plate (300 kV). The hole in the phase plate was 15 μm wide, which corresponds to 0.11/Å cut-on frequency. The unscattered beam was centered in the hole. The DQE curve fall-off is the same as for a VPP without a hole.

**Figure 8.**
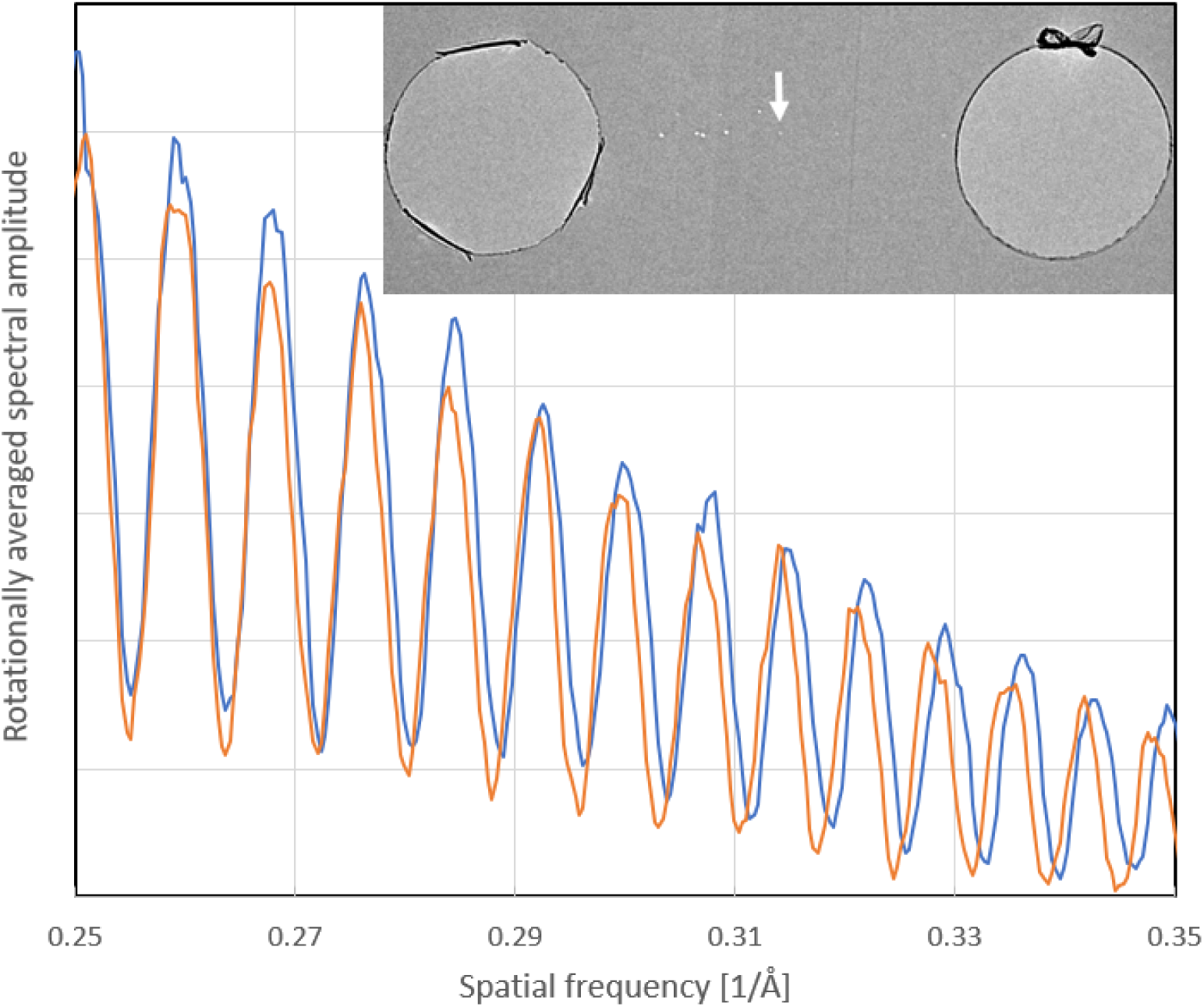
Rotationally averaged Thon rings of thin amorphous carbon obtained with a ∼3nm thin phase plate (300 kV). The inserted figure shows the phase plate layout with two holes that have a diameter of 15 μm. The unscattered beam was focused on the carbon film between these two holes at the position of the white arrow. The rotational average of the PSD of the image of amorphous carbon is taken in an azimuthal section that covers the hole (blue curve) and a similar-sized azimuthal section outside the hole (orange curve). The modulation depth is the same, within experimental error. The blue curve has somewhat higher overall intensity because there is less scattering by the phase plate (as the electron wave is passing the two holes).

## DISCUSSION

We show here that the signal-loss associated with the VPP is not constant as a function of spatial frequency. Instead, we observe a progressive reduction in signal with increasing resolution. At ∼2 Å resolution, for example, the high-resolution signal is reduced to as little as ∼40% of that observed without a VPP. This reduction effectively represents a resolution-dependent decrease in the detective quantum efficiency, DQE(s), which is attributable solely to the VPP.

The unwanted loss of high-resolution signal clearly acts in opposition to the benefits expected from using the VPP for imaging smaller biological macromolecules. This previously-unexpected loss of high-resolution signal may thus be a reason why neither the resolution achieved, nor the number of particles required to obtain a high-resolution reconstruction, have been shown to be improved when a VPP is used for single-particle cryo-EM. On the contrary, the results achieved for specimens such as hemoglobin (Herzik et al., 2019) or GPCR membrane proteins (Zhang et al., 2017) are almost indistinguishable from those achieved previously with a VPP (Khoshouei et al., 2017, Liang et al., 2017). Such comparisons are difficult to make, however, especially if the differences are small. This is because many other factors may be involved, such as differing ice thicknesses, differing degrees of structural heterogeneity, or differing procedures used for data processing.

Another reason why VPP performance at high resolution can be compromised is the fact that variations in VPP surface topography (wrinkling) cause aberrations in the electron wave passing the VPP. Aberrations beyond 2-fold astigmatism are difficult to correct with CTF correction software. Recent tests with flattened VPP films have shown that these aberrations can be reduced significantly. Future testing will show if the resolution limitation is dominated by DQE fall-off at high resolution or by VPP-induced optical aberrations.

On the other hand, the fall-off of signal at high resolution, which we observe here, does not significantly compromise the value added by using the VPP for most applications in biological electron cryo-tomography. This is because the DQE of the VPP remains quite high at the resolutions normally achieved in tomography, which – for other reasons – are rarely as high as ∼1 nm. On the contrary, there is a significant benefit to using defocus values as small as 1 µm or less for cryo-tomography, which is possible when using a phase plate. Indeed, the image quality achieved with a VPP exceeds that achieved by just using an energy filter, although it is even better to use both (Fukuda et al., 2015).

A number of experiments were performed in an effort to better understand the unexpected signal loss at high resolution. Hypotheses that might account for the unwanted DQE(s) can be grouped into two possible categories, corresponding to either surface effects or bulk effects in the material of the phase plate. We believe that two key results are difficult to explain by known interactions of electrons with bulk materials. They are: 1) The fall-off of DQE is independent of the thickness of the VPP, to within experimental error. This is in spite of the fact that the signal does decrease measurably, but to the same extent at all resolutions, with increasing thickness of the carbon film. 2) The amount of fall-off is hardly sensitive to whether the electron energy is 200 keV or 300 keV. The fall-off of the DQE of the VPP thus appears more likely to be caused by a surface effect, just as the Volta potential itself is thought to be. One possible suggestion might still be that rapid spatial fluctuations in surface charge of the carbon film, such as those believed to cause the “bee swarm” effect (Russo and Henderson, 2018, Dove, 1964), act as a fluctuating beam-diffuser. If such beam diffusion causes the observed DQE effect, it must happen for both the unscattered and the scattered part of the electron beam, given the fact that even a hole-based phase plate shows a similar DQE fall-off.

## ACKNOWLEDGMENTS

We acknowledge support in phase plate manufacturing and installation by Erwin van Hoek, Richard Bolt, and Roland Jonkers. This work was supported in part by the US National Institutes of Health grant 5 R01 GM126011, and Japan Science and Technology Agency (JST) PRESTO grant #18069571.

